# IscR-mediated sensing of iron-sulfur cluster demand coordinates virulence gene expression in *Yersinia pseudotuberculosis*

**DOI:** 10.1101/2025.10.16.682990

**Authors:** Mané Ohanyan, George Imre Balazs, Karthik Hullahalli, Diana Fernandez Pacheco, Caia Lomeli, Katherine G. Dailey, Erin Mettert, Diana Hooker-Romero, Leah Schwiesow, Patricia J. Kiley, Matthew K. Waldor, Victoria Auerbuch

## Abstract

The type III secretion system (T3SS) is a needle-like appendage that translocates effector proteins into host cells to disrupt host defenses. Strict control of T3SS expression is critical for facultative pathogens such as *Yersinia pseudotuberculosis*, as the T3SS is indispensable for virulence but is metabolically costly. We previously showed that the iron-sulfur (Fe-S) cluster-coordinating transcription factor IscR controls expression of LcrF, the master regulator of the *Yersinia* T3SS and the YadA adhesin. Clusterless apo-IscR, the predominant IscR form during high cellular Fe-S cluster demand (aerobic, low iron conditions), promotes LcrF, T3SS, and YadA expression. Importantly, binding of apo-IscR to the *lcrF* promoter at a type II IscR binding site facilitates *Yersinia* disseminated infection. Here, we show that mutating the *lcrF* promoter to allow only [2Fe-2S]-IscR binding (*lcrF*^pTypeI^) results in hyperexpression of LcrF, the T3SS, and YadA during low Fe-S cluster demand (anaerobic, iron replete conditions). These data suggest that switching the form of IscR that can bind to the *lcrF* promoter reversed how iron and oxygen regulate *Yersinia* virulence factors. We used barcoded *Y. pseudotuberculosis* to probe how control of the *lcrF* promoter in response to iron and oxygen modifies infection dynamics. We found that the *lcrF*^pTypeI^ mutant experiences a tighter bottleneck in the cecum, where *Yersinia* is expected to experience a low oxygen, iron replete environment. Taken together, these findings suggest that by tying T3SS and YadA expression to cellular Fe–S cluster demand, *Yersinia* can fine-tune its virulence repertoire to the host tissue microenvironment.

**Importance:** Iron and oxygen availability fluctuate spatially across mammalian tissues as well as temporally during the course of bacterial infection. The [2Fe-2S] cluster coordinating transcription factor IscR senses changes in iron and oxygen levels, and plays a pivotal role in enabling pathogens like *Yersinia*, *Salmonella*, and *Vibrio* to express critical virulence genes. While prior research has established that iron availability and oxygen tension influence IscR abundance and DNA-binding specificity, it is unclear how these changes control the timing and location of virulence factor expression during infection. In this study, we engineered a bacterial strain to reverse the way in which iron and oxygen drive expression of two critical virulence factors through IscR. This mutant displayed altered host infection dynamics, revealing that uncoupling virulence gene expression from host tissue cues decreases bacterial fitness.

## Introduction

*Yersinia pseudotuberculosis* is a Gram-negative, food-borne pathogen that causes mesenteric lymphadenitis in otherwise healthy human hosts, and more serious disseminated infection in individuals with iron overload disorders such as hereditary hemochromatosis^1,2^. *Y. pseudotuberculosis* serves as a model species for studying enteric infections in mammalian hosts because oral infection of mice does not require antibiotic pre-treatment and leads to colonization of intestinal, lymph, and vital organ tissue^3^. *Y. pseudotuberculosis* as well as the closely related plague agent *Y. pestis* carry a ∼70 kb virulence plasmid (<u>p</u>lasmid of *<u>Y</u>ersinia* <u>v</u>irulence, <u>pYV</u>) that encodes the Ysc type III secretion system (T3SS)^4,5^. The T3SS is a needle-like appendage that injects *<u>Y</u>ersinia* <u>o</u>uter <u>p</u>roteins (<u>Yops</u>) into target host cells to interfere with phagocytosis, inhibit cytokine and reactive oxygen species production, and modulate other host defense mechanisms^6,7^. Importantly, the T3SS is a conserved virulence mechanism across many other clinically significant pathogens including *Salmonella, Escherichia, Shigella, Vibrio, Pseudomonas* and more. While the T3SS is central to the ability of these pathogens to infect mammalian tissue, T3SS expression is metabolically burdensome. The T3SS contains antigens recognized by the innate and adaptive immune responses, and in some cases is only required for bacterial survival in specific host tissue niches ^8,9,10,11,12^. Indeed, *in vitro* T3SS activity is associated with growth arrest in *Yersinia*, *Salmonella*, and *Shigella*^13,14^. Not surprisingly, expression of the T3SS is tightly regulated. To balance the virulence requirement of the T3SS with its fitness burden, *Y. pseudotuberculosis* regulates its T3SS at the level of (i) pYV copy number and therefore T3SS gene dosage, (ii) transcription, mRNA decay, and translation of T3SS genes, and (iii) T3SS activity^15,16,17^. The master transcriptional regulator of the Ysc T3SS is the pYV-encoded, AraC family transcription factor LcrF^18,19,20^. LcrF is only translated at mammalian body temperature as a result of an RNA thermometer and positively regulates the T3SS structural components, chaperones, and Yops necessary for assembly and function of the T3SS^21, 22^.

Several regulatory mechanisms that control *lcrF* transcription have been described. The *lcrF* gene is part of a two-gene operon, *yscW-lcrF*, with the upstream gene encoding the T3SS pilotin protein YscW^23^. The *yscW-lcrF* promoter is repressed by the H-NS global repressor of horizontally acquired genes and its binding partner YmoA^24^, and activated by the iron-<u>s</u>ulfur <u>c</u>luster <u>r</u>egulator IscR^25,26^. IscR is a global transcription factor that regulates the Isc and Suf iron sulfur cluster biogenesis pathways responsible for forming iron-sulfur (Fe-S) clusters for target apo-proteins^27^. IscR itself is an Fe-S cluster coordinating protein, capable of both activating and repressing gene expression in its clusterless apo-IscR form, and its [2Fe-2S] cluster- bound holo-IscR form^28, 29, 30, 31^. Apo-IscR binds target DNA sequences known as type 2 motifs ^29, 32, 31^. While holo-IscR binds both type 1 and type 2 motifs, holo-IscR binding to promoters containing type 2 motifs may not impact transcription in the same way as apo-IscR binding of type 2 motifs^33^. Cellular ratios of holo-IscR and apo-IscR are controlled by cellular Fe-S cluster demand^34, 32^. This is primarily because holo-IscR binds two tandem type 1 motifs in the *iscRSUA-hscBA-fdx-iscX* operon and represses expression of IscR and the Isc Fe-S cluster biogenesis system^28,34^. This results in overall low levels of cellular IscR, with the predominant form being holo-IscR. However, conditions like iron starvation or oxidative stress are known to increase demand for Fe-S cluster biogenesis or damage Fe-S clusters, respectively ^35,36,37^. Therefore, in these conditions holo-IscR levels decrease, relieving the repression on the *iscRSUA-hscBA-fdx-iscX* operon and increasing overall IscR levels, predominantly in the apo-IscR form^27^. In *Yersinia*, IscR binds to a type 2 motif in the *yscW-lcrF* promoter that is flanked by two H-NS binding sites^25,24^. IscR is required for *yscW-lcrF* expression specifically in the presence of H-NS/YmoA^24^, suggesting that IscR prevents H-NS/YmoA repression of the *yscW-lcrF* promoter. This appears to occur specifically under iron limiting or aerobic conditions when Fe-S cluster demand is high and apo-IscR predominates^26^.

A mouse oral infection model revealed that a *Y. pseudotuberculosis* Δ*iscR* mutant is defective in colonizing the Peyer’s patches, small intestine, spleen, and liver ^25^. However, IscR directly regulates dozens of genes in *Yersinia*^38^. A more discrete mutation of the type 2 motif in the *yscW-lcrF* promoter led to a *Y. pseudotuberculosis* strain that expresses IscR, but lacks IscR binding to the *yscW-lcrF* promoter (*lcrF*^pNull^)^26^. This strain colonized the small intestine lumen normally, but was defective in disseminated infection, suggesting that IscR-dependent upregulation of the T3SS is dispensable in the intestine, but is required for *Yersinia* virulence in extraintestinal tissues where the T3SS is needed to counter innate immune defenses. Interestingly, *Yersinia* is known to experience anaerobic, iron replete conditions in the intestinal lumen ^39, 40, 41, 42, 43, 44^, which should lead to low Fe-S cluster demand and overall low levels of mostly holo-IscR. In contrast, *Yersinia* experiences iron starvation in extraintestinal tissues ^45^, which should lead to higher Fe-S cluster demand and higher levels of mostly apo-IscR. However, it remains unclear when IscR sensing of iron and oxygen becomes important during *Yersinia* infection of a mammalian host.

To probe how IscR sensing of iron and oxygen regulates T3SS expression dynamics and *Yersinia* virulence, we swapped the previously characterized native type 2 IscR DNA binding motif in the *Y. pseudotuberculosis yscW-lcrF* promoter to a type 1 motif (*lcrF*^pType1^). We found that, while wildtype bacteria expressed the T3SS under aerobic or iron limiting conditions, the *lcrF*^pType1^ mutant expressed the T3SS preferentially under anaerobic iron replete conditions. As a consequence, the *lcrF*^pType1^ mutant experienced a tighter bottleneck and was defective in colonizing the cecum where the *lcrF*^pType1^ mutant, but not wildtype *Y. pseudotuberculosis*, is predicted to express the T3SS. These data provide insight into how bacteria use IscR to sense their biogeography to control virulence factor expression.

## Results

### The *lcrF*^pType1^ mutant exhibits increased T3SS activity relative to WT under anaerobic iron replete conditions

To investigate how IscR contributes to *lcrF* regulation in response to environmental cues, we replaced the IscR type 2 binding motif in the *yscW*-*lcrF* promoter with a type 1 motif in a β-galactosidase reporter construct (*lcrF*^pType1^ mutation, Fig 1A). As expected, apo-IscR bound to the wildtype (WT) but not the *lcrF*^pType1^ mutant *yscW-lcrF* promoter (Fig S1A). In contrast, holo-IscR retained the ability to bind to the *lcrF*^pType1^ mutant promoter (Fig S1B). We hypothesized that conditions favoring holo-IscR will induce activity of the *lcrF*^pType1^ mutant promoter but not the WT promoter. Indeed, we found that the pType1 mutation led to significantly higher *yscW-lcrF* promoter activity than the native type 2 motif under anaerobic iron replete conditions (Fig 1B). In contrast, both reporters exhibited similarly low levels of promoter activity in a Δ*iscR* mutant background, suggesting that the reporter activity difference observed in the WT strain was IscR-dependent.

**Figure 1.**
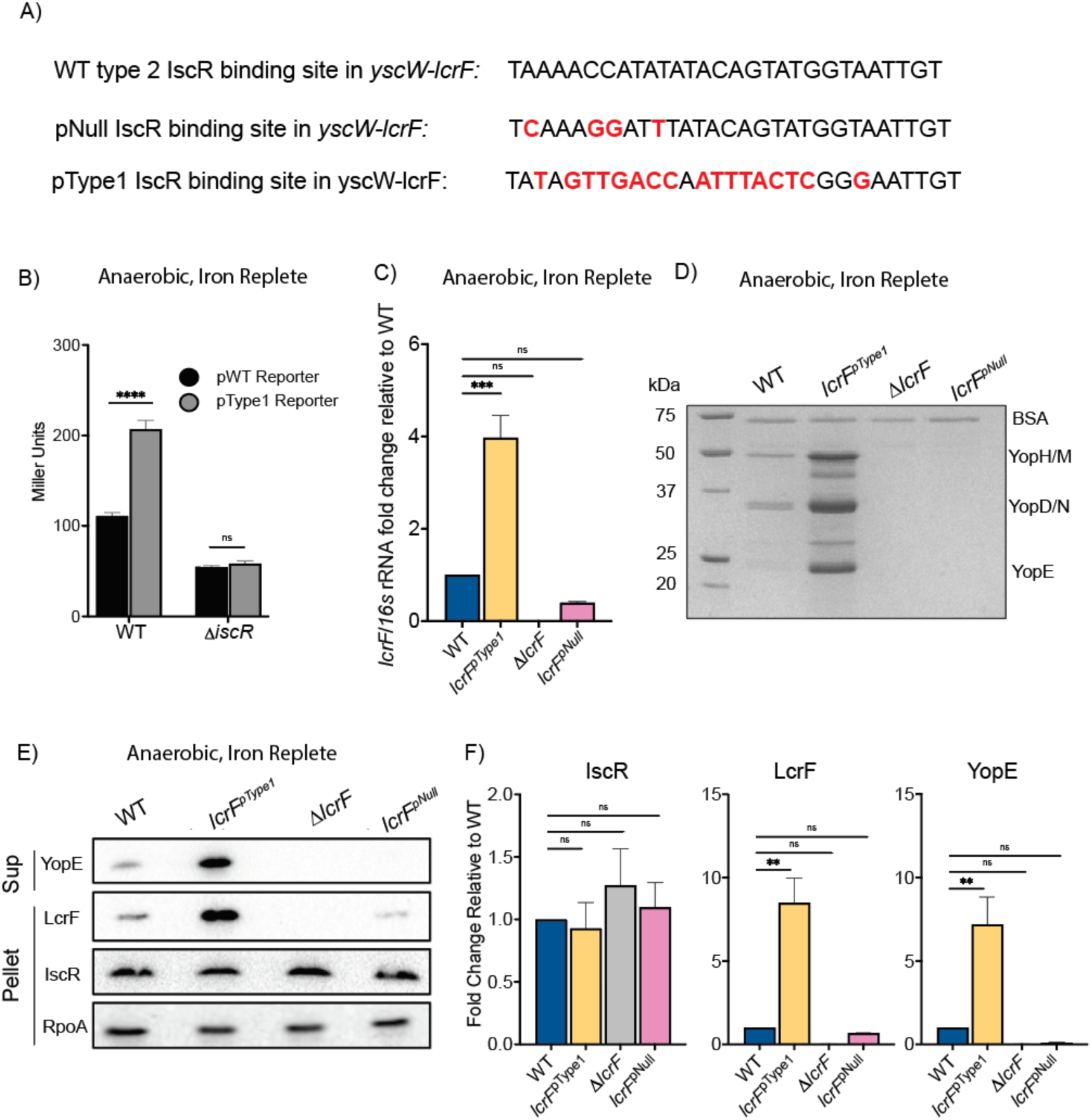
The *lcrF*^pType1^mutation leads to hyperactivity of the T3SS under anaerobic iron replete conditions. **A)** IscR binding site sequences in the *yscW-lcrF* promoter. Base pair substitutions to generate the pType1 and pNull mutations indicated in red. **B-F)** All strains were cultured anaerobically for 1 hour at room temperature then switched to 37° C for 5 hours for continued anaerobic incubation. **B)** β-galactosidase assays were carried out with WT and Δ*iscR Y. pseudotuberculosis* harboring pFU99-encoded p*yscW-lcrF* transcriptional reporters containing either the WT type 2 motif or pType 1 mutant promoters. Promoter activity measured in Miller Units. The average ± SEM of at least three independent experiments is shown. Two-way ANOVA Bonferroni’s multiple comparisons test. ****p<0.0001, ns, not significant. **C)** Quantitative RT-PCR was used to measure *lcrF* transcript levels in the WT, Δ*lcrF*, *lcrF*^pNull^, and *lcrF*^pType1^ strains relative to 16s rRNA. Bar graph shows the average fold change relative to WT for three independent experiments ± SEM. One-way ANOVA Tukey’s multiple comparisons test. **** p<0.0001, ns, not significant. **D)** Proteins secreted into the culture supernatant and precipitated with trichloroacetic acid were separated by SDS-PAGE and visualized with Coomassie blue. **E)** Representative western blot showing relative pellet LcrF, IscR, and RpoA as well as supernatant (sup) YopE protein levels. **F)** Average relative quantification of western blot data from at least three independent experiments ± SEM is shown. One-way ANOVA with Tukey’s multiple comparisons test ****p<0.0001, ns, not significant.

To characterize how the pType 1 mutation in the *yscW-lcrF* promoter impacts T3SS activity, a *Y. pseudotuberculosis lcrF*^pType1^ strain was made by replacing the type 2 motif in the native *yscW-lcrF* promoter with the pType 1 sequence. Two additional strains were used as negative controls, Δ*lcrF* and *lcrF*^pNull^, which either lack the *lcrF* gene or have a mutation in the *yscW-lcrF* promoter that abolishes IscR binding and prevents *lcrF* transcription, respectively^26^. Under anaerobic iron replete conditions that promote low cellular Fe-S cluster demand and favor holo-IscR, *lcrF* mRNA levels were not significantly different between the WT, Δ*lcrF*, and *lcrF*^pNull^ strains. In contrast, *lcrF* transcripts were increased four-fold in the *lcrF*^pType1^ mutant relative to the WT strain (Fig 1C). Furthermore, analysis of secreted effectors in the culture supernatant showed T3SS hyperactivity in the *lcrF*^pType1^ mutant relative to WT under these conditions (Fig 1D). A similar pattern was observed for LcrF protein levels as well as levels of the T3SS effector protein YopE, an LcrF target gene (Fig 1E,F). IscR protein levels were similar across all strains, suggesting that the difference observed in LcrF expression and activity in the *lcrF*^pType1^ strain likely results from holo-IscR activity on the *yscW-lcrF* promoter in the *lcrF*^pType1^ strain but not the WT strain.

### The *lcrF*^pType1^ mutant exhibits decreased T3SS activity relative to WT in aerobic iron limiting conditions

In aerobic iron limiting conditions that promote high cellular Fe-S cluster demand, the predominant form of IscR is predicted to be apo-IscR, which is unable to bind type 1 motif sequences^31,27^. We hypothesized that while the *lcrF*^pType1^ mutant exhibited a hypersecretion phenotype in anaerobic, iron replete conditions that favor holo-IscR, aerobic iron limiting conditions that drive IscR toward the apo- form should render the *lcrF*^pType1^ mutant defective in T3SS expression relative to WT. To test this hypothesis, we starved *Y. pseudotuberculosis* strains of iron and grew them in the presence of oxygen. Using the same β-galactosidase reporters as described earlier, the activity of the *lcrF*^pType1^ mutant promoter was significantly less than in the WT promoter (Fig 2A). Importantly, there was no difference in the promoter activity of the two reporters in the Δ*iscR Y. pseudotuberculosis* strain, suggesting that the differences observed in the WT *Y. pseudotuberculosis* background were IscR- dependent. Consistent with our prediction, the *lcrF*^pType1^ *Y. pseudotuberculosis* strain exhibited significantly lower *lcrF* mRNA and LcrF protein levels, as well as YopE secretion, relative to WT (Fig 2B-E). The *lcrF*^pType1^ mutant had slightly greater abundance of *lcrF* mRNA and YopE expression than the *lcrF*^pNull^ mutant, suggesting that some holo-IscR is present under these conditions to drive a small amount of T3SS expression. Taken together, these data suggest that swapping the native *yscW-lcrF* promoter type 2 motif with a type 1 motif that can only be bound by holo-IscR leads to decreased LcrF and T3SS expression under conditions where high Fe-S cluster demand keeps holo-IscR levels low.

**Figure 2.**
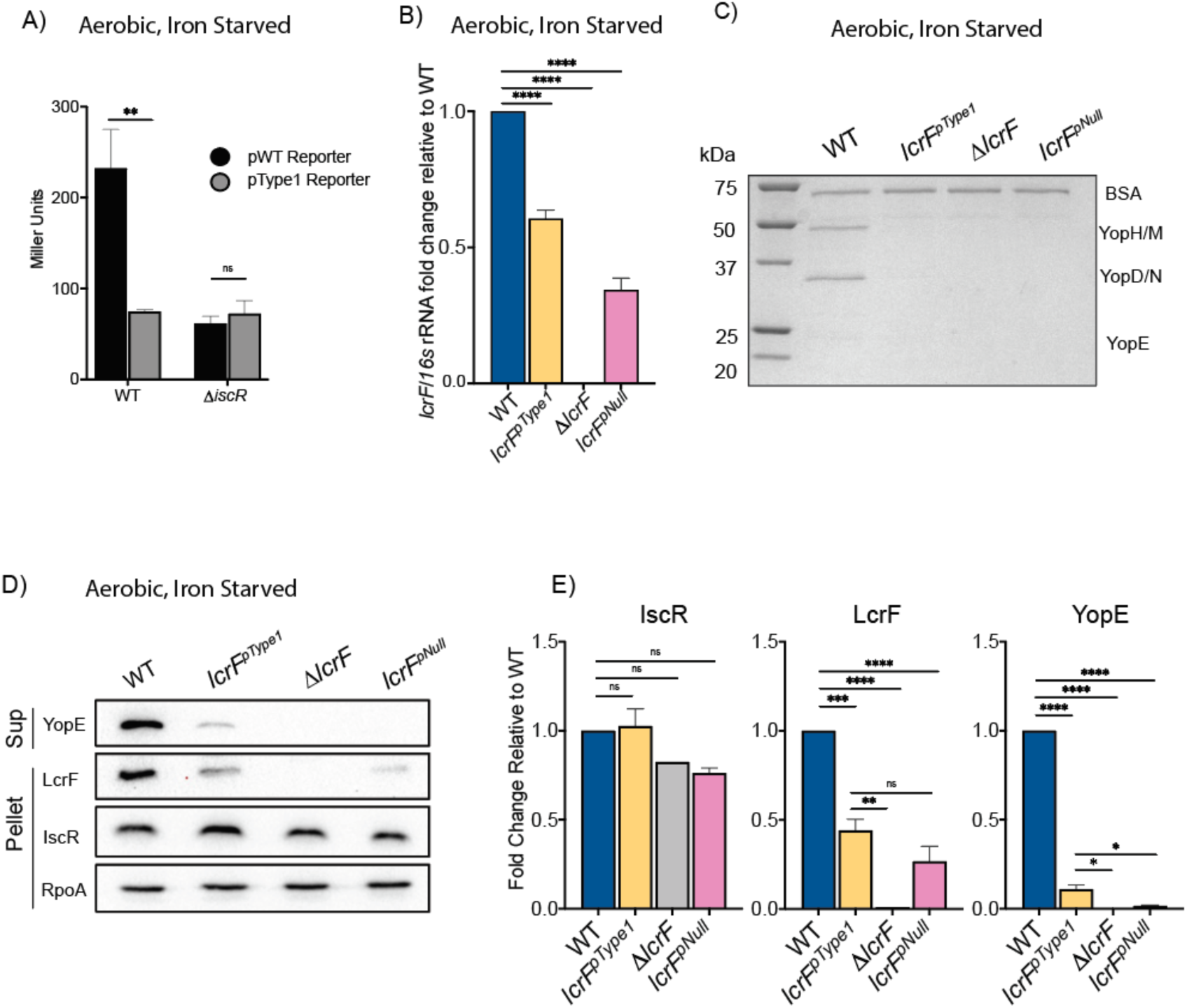
*Y. pseudotuberculosis* leverages its native type 2 IscR binding motif in the *yscW-lcrF* promoter to facilitate T3SS activity under aerobic iron limiting conditions. All strains were cultured aerobically and starved for iron at room temperature then switched to 37° C for 4 hours for continued aerobic incubation. **A)** β-galactosidase assays were carried out with WT and Δ*iscR Y. pseudotuberculosis* harboring pFU99-encoded p*yscW-lcrF* transcriptional reporters containing either the WT type 2 motif or pType 1 mutant promoters. Promoter activity measured in Miller Units. The average ± SEM of at least three independent experiments is shown. Two-way ANOVA Bonferroni’s multiple comparisons test. ****p<0.0001, ns, not significant. **B)** Quantitative RT-PCR was used to measure *lcrF* transcript levels in the WT, Δ*lcrF*, *lcrF*^pNull^, and *lcrF*^pType1^ strains relative to 16s rRNA. Bar graph shows the average fold change relative to WT for three independent experiments ± SEM. One-way ANOVA Tukey’s multiple comparisons test. **** p<0.0001, ns, not significant. **C)** Proteins secreted into the culture supernatant and precipitated with trichloroacetic acid were separated by SDS-PAGE and visualized with Coomassie blue. **D)** Representative western blot showing relative pellet LcrF, IscR, and RpoA as well as supernatant (sup) YopE protein levels. E) Average relative quantification of western blot data from at least three independent experiments ± SEM is shown. One-way ANOVA with Tukey’s multiple comparisons test ****p<0.0001, ns, not significant .

### The *lcrF*^pType1^ mutant exhibits altered expression of the LcrF target gene *yadA* in response to changes in iron and oxygen availability

YadA is an LcrF-regulated virulence factor encoded on pYV. Trimeric YadA complexes are expressed on the surface of *Yersinia* and have been implicated in several important physiological processes such as β1 integrin-mediated binding to host cells and serum resistance ^46^. In our effort to elucidate the broader relevance of IscR binding dynamics at the *yscW-lcrF* promoter, we tested the impact of the pType1 mutation in the *yscW-lcrF* promoter on *yadA* expression levels in either anaerobic iron replete media or aerobic iron limiting media. As expected, in anaerobic iron replete media the pType1 mutation leads to a two-fold increase in *yadA* mRNA levels relative to WT (Fig 3A). Furthermore, we observed the inverse phenotype in aerobic iron limiting conditions, where the *lcrF*^pType1^ strain exhibited half the *yadA* mRNA levels observed in WT (Fig 3B). Taken together, our data suggest that the way in which iron availability and oxygen tension control expression of the *Yersinia* T3SS and YadA can be switched by changing the type of IscR binding site present in the *yscW-lcrF* promoter, and therefore which form of IscR can control *yscW-lcrF* transcription.

**Figure 3:**
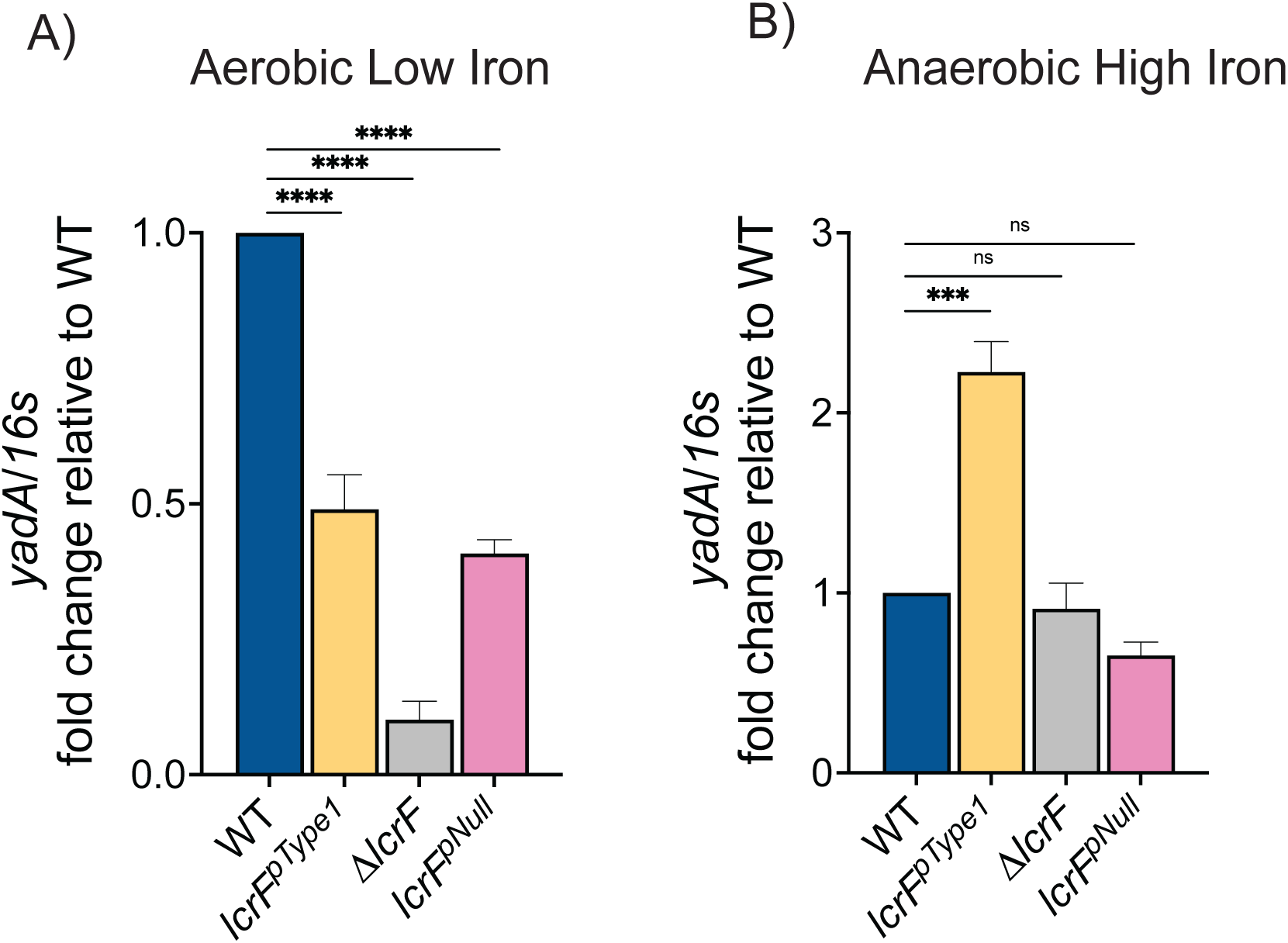
Swapping the form of IscR that controls the *yscW-lcrF* promoter reverses how iron and oxygen regulate YadA expression. A-B) Quantitative RT-PCR was used to measure *lcrF* transcript levels in the WT, Δ*lcrF*, *lcrF*^pNull^, and *lcrF*^pType1^ strains relative to 16s rRNA. Bar graph shows the average fold change relative to WT for three independent experiments ± SEM. One-way ANOVA Tukey’s multiple comparisons test. **** p<0.0001, ns, not significant. **A)** Strains were cultured anaerobically for 1 hour at room temperature then switched to 37° C for 5 hours for continued anaerobic incubation as in Figure 1. **B)** Strains were cultured aerobically and starved for iron at room temperature then switched to 37° C for 4 hours for continued aerobic incubation as in Figure 2.

### Changing only oxygen tension while maintaining iron repletion leads to misregulation of the T3SS in the *lcrF*^pType1^ mutant

We grew *Y. pseudotuberculosis* in non-chelated iron replete media under either aerobic or anaerobic conditions for a direct comparison of *Yersinia* under high and low Fe-S cluster demand, respectively. As expected, IscR levels were significantly elevated in the presence of oxygen compared to the absence of oxygen (Fig 4A,B). These findings support the model that oxidative stress increases Fe-S cluster demand leading to a large pool of mostly apo-IscR, whereas anaerobic conditions facilitate low Fe-S cluster demand that promotes a smaller pool of mostly holo-IscR. In line with our previous data, LcrF levels were significantly higher in aerobic compared to anaerobic conditions for the WT strain (Fig 4A, C). LcrF expression in WT was nearly double that observed in the *lcrF*^pType1^ strain under aerobic conditions, and about four times more than *lcrF*^pNull^ (Fig 4A, Fig 4C). Conversely, when cultured anaerobically, the *lcrF*^pType1^ mutant exhibited approximately 5-fold more LcrF expression relative to WT. Interestingly, comparing LcrF expression in the *lcrF*^pType1^ strain in the presence or absence of oxygen showed only a slight, non-significant increase in the anaerobic condition. This suggests the presence of holo-IscR in oxygenated iron-replete media, and is consistent with our previous results^25^. Taken together, these data suggest that the native IscR type 2 binding motif in the *lcrF* promoter enables *Y. pseudotuberculosis* to preferentially express the T3SS in aerobic environments, whereas placing *lcrF* under the control of a type 1 motif eliminates the ability of *Yersinia* to prevent T3SS expression anaerobically.

**Figure 4:**
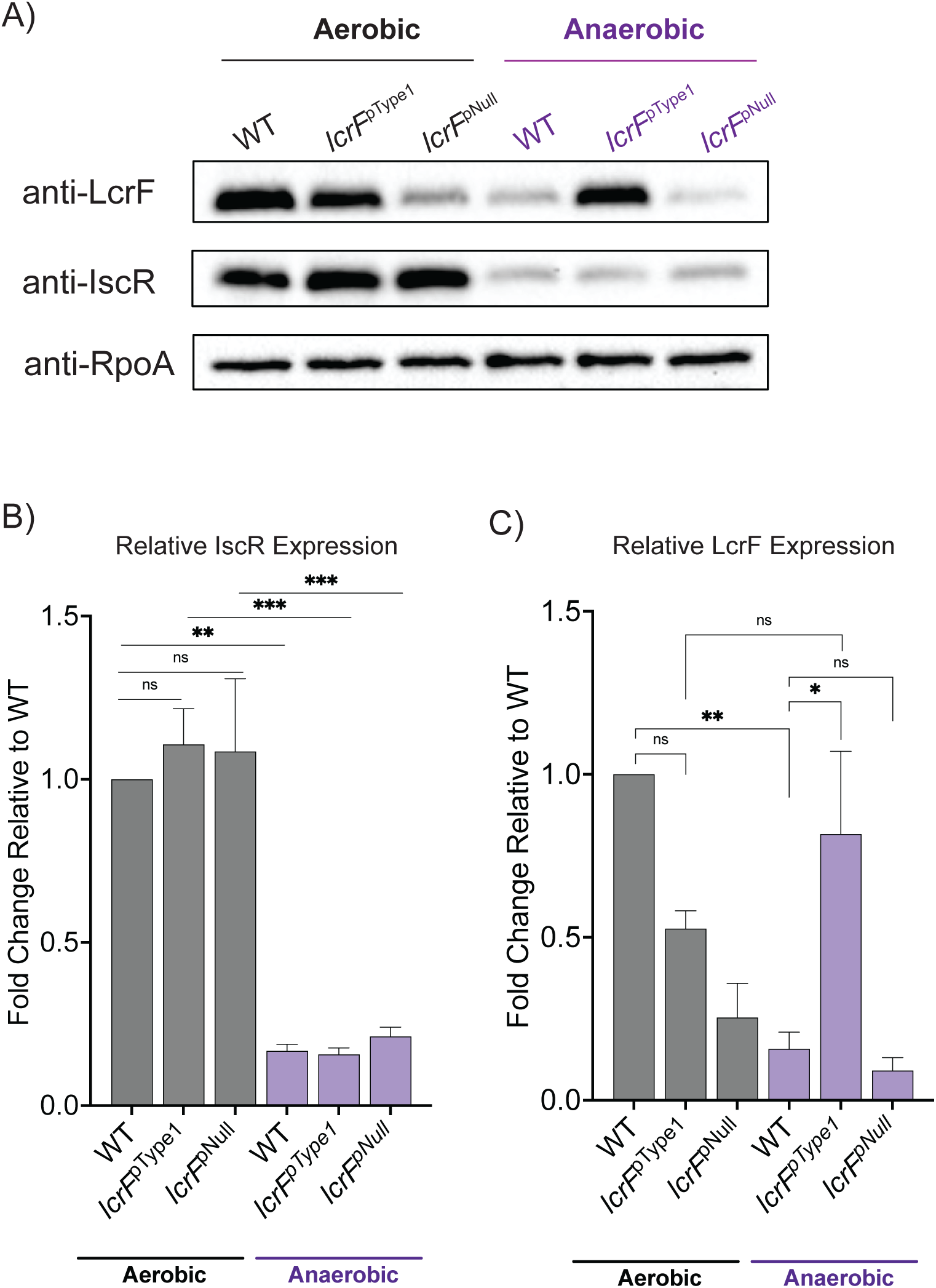
The IscR type 2 motif in the *lcrF* promoter enables *Yersinia* to express the T3SS aerobically while preventing T3SS expression anaerobically. Strains were cultured in M9, non-chelated media and incubated either aerobically or anaerobically to assess T3SS activity. **A)** Western blot showing LcrF, IscR, and RpoA protein levels in the pellet. **B)** Average fold change in IscR levels normalized to RpoA and relative to the aerobic WT condition for three independent experiments ± SEM. **C)** Average fold change in LcrF levels normalized to RpoA and relative to the aerobic WT condition for three independent experiments ± SEM. Statistical significance determined by one-way ANOVA Bonferroni’s multiple comparisons test. *p<0.05; **p<0.01; ***p<0.001; ns, not significant.

### IscR mediated control of the *yscW-lcrF* promoter is important for cecal colonization

To test how mis-regulation of LcrF, the T3SS, and YadA in response to iron and oxygen affects *Yersinia* virulence, employed a mouse oral feeding model. Mice infected with WT *Y. pseudotuberculosis* trended toward losing more weight during the course of infection compared to mice infected with the *lcrF*^pType1^ mutant (Fig 5A), although this difference was not statistically significant. Similarly, fecal myeloperoxidase (MPO), an enzyme released by activated neutrophils, trended lower in *lcrF*^pType1^-infected mice, although this was not statistically significant (Fig 5B).

**Figure 5:**
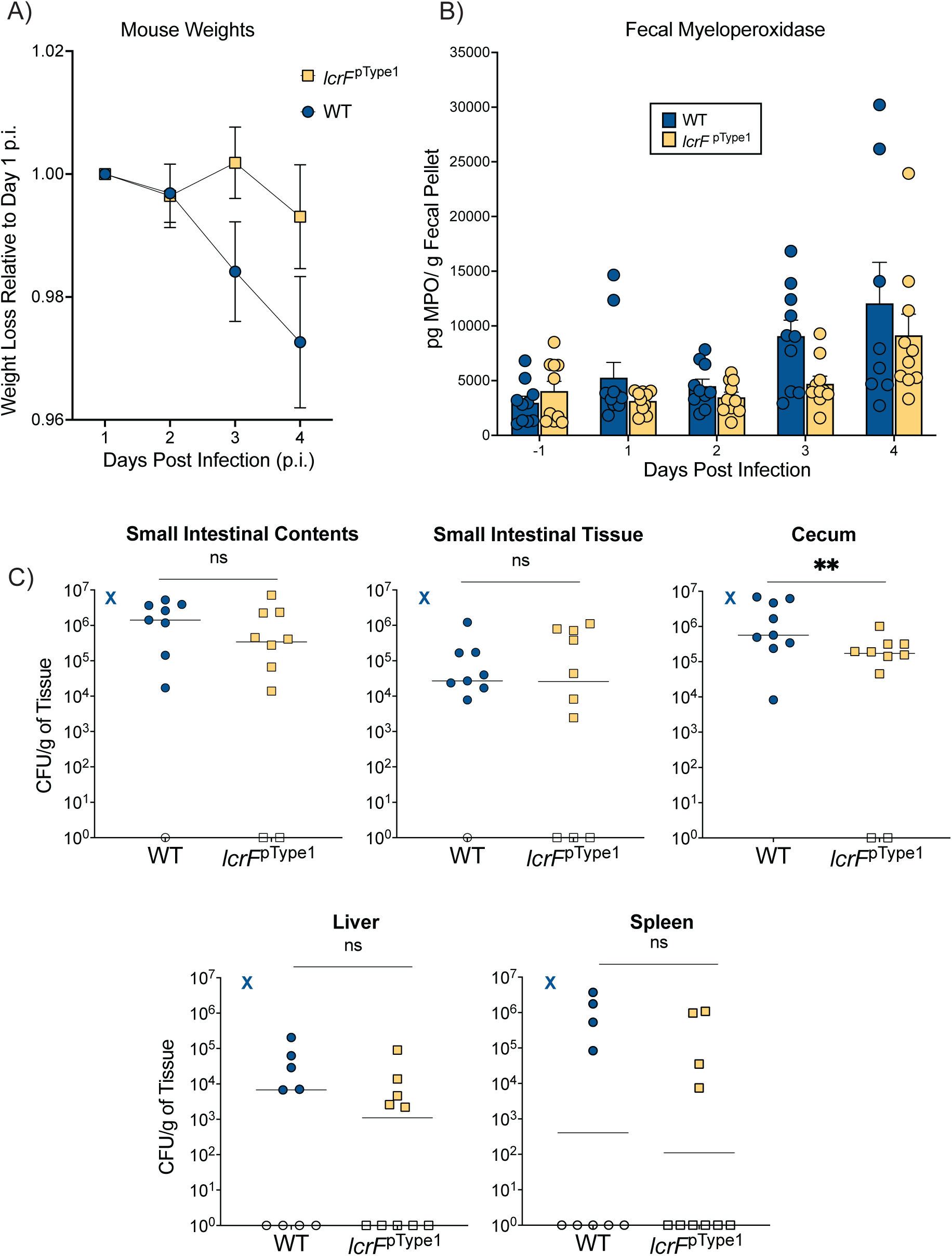
Swapping the form of IscR that controls the *lcrF* promoter disrupts cecal colonization. C57BL/6 mice were infected via feeding with 10^9^ CFU of *Y. pseudotuberculosis* WT or *lcrF*^pType1^. **A)** Daily weight for each mouse was measured relative to its weight at day 1 post-inoculation. The average ± SEM of 10 mice is shown. **B)** Fecal samples were collected one day prior, or 1-4 days post-inoculation and myeloperoxidase (MPO) levels measured via ELISA. Each symbol presents data from one mouse. Bars show average ± SEM. **C)** Bacterial burden in small intestinal contents and tissue, cecum, spleen, and liver were determined four days post- inoculation. Each circle or square represents data from one mouse. Open symbols indicate no CFU detected. Lines represent geometric mean. “X” indicates a mouse that exhibited signs of severe illness and was therefore euthanized prior to the end of the experiment. Statistical significance as determined by unpaired Mann-Whitney test. **p<0.01; ns is not significant. Data is from two independent experiments.

However, cecal bacterial burden was significantly lower in the *lcrF*^pType1^-infected mice compared to WT four days post-inoculation (Fig 5C). Interestingly, the cecal lumen is predicted to be an anaerobic environment, which we have shown *in vitro* induces T3SS and YadA expression in the *lcrF*^pType1^ mutant but not in WT *Y. pseudotuberculosis*.

### Misregulation of the *yscW-lcrF* promoter tightens the bottleneck to cecal colonization

To better understand how *yscW-lcrF* misregulation impacts *in vivo* infection dynamics of *Y. pseudotuberculosis*, we employed Sequence Tag-based Analysis of Microbial Populations in R (STAMPR). In this method, barcoded but otherwise isogenic bacteria are used to quantify infection bottlenecks^47^. We generated barcoded libraries in WT and *lcrF*^pType1^ *Y. pseudotuberculosis* strain backgrounds by integrating ∼60,000 barcodes of ∼25 nucleotides in length into a neutral site on the chromosome. *In vitro* bottlenecks created by serial dilutions were used to validate that these libraries could be used to measure founding populations ranging from 1^0^ to 10^5^ using the STAMPR analysis pipeline (Fig S2). The founding population, defined via a metric known as Ns, defines the number of cells from the inoculum that give rise to the bacterial population at the site of infection. Smaller founding population sizes relative to the inoculum size indicate tighter bottlenecks to colonization, where only a small number of clones can establish infection. In contrast, larger values suggest that many unique bacterial clones were able to seed and colonize a given organ, indicative of a wider bottleneck. We found that in the cecum of mice infected with the *lcrF*^pType1^ mutant, the Ns value was significantly less than that of WT, suggesting that *yscW-lcrF* misregulation tightens the cecal colonization bottleneck (Fig 6). While Ns values between the two strains in the other organs were not statistically different, it is striking that mice infected with *lcrF*^pType1^ had significantly smaller Ns values in the spleen and the liver relative to WT, suggesting a possible tighter bottleneck to disseminated infection. Taken together, these data indicate that the ability of *Y. pseudotuberculosis* to express T3SS and YadA specifically in response to low iron and oxygen enables strategic virulence factor deployment and promotes tissue colonization. Furthermore, these data suggest that expressing the T3SS and YadA in response to anaerobic iron replete conditions may be detrimental to *Y. pseudotuberculosis* virulence.

**Figure 6.**
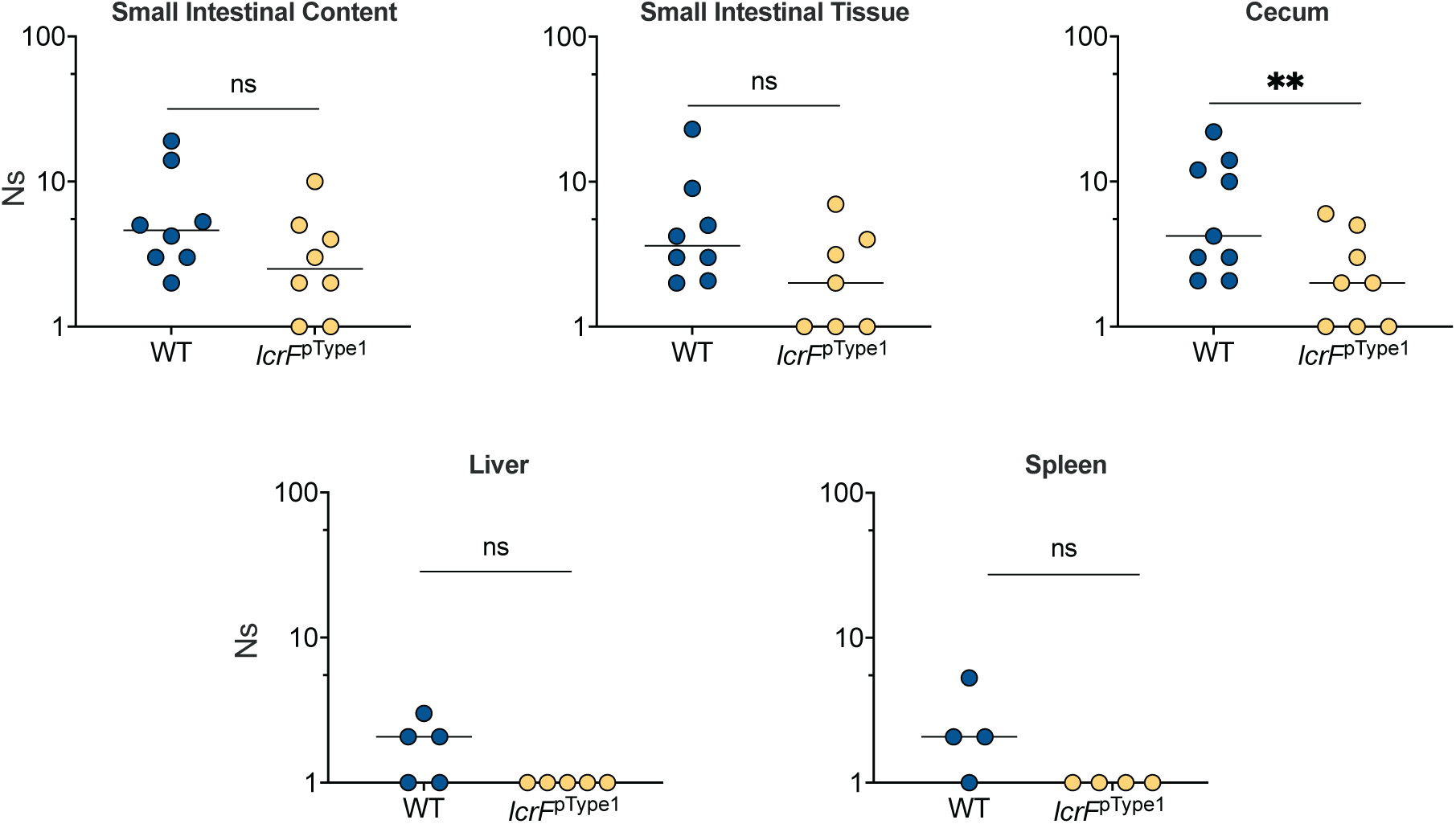
Misregulation of the *yscW-lcrF* promoter tightens infection bottlenecks. Founding populations (Ns) for each organ shown in Fig 5 were determined by comparing barcode frequencies in the undiluted STAMPR library to those at each tissue site, using the STAMPR pipeline Statistical significance as determined by unpaired Mann-Whitney test, **p<0.01, ns is not significant.

## Discussion

In this study, we developed a *Y. pseudotuberculosis* mutant strain in which the native IscR type 2 binding motif in the promoter of the LcrF T3SS master regulator is swapped with a type 1 motif (*lcrF*^pType1^). This mutation effectively exchanges the form of IscR that controls the *lcrF* promoter, from apo-IscR to holo-IscR (Fig 7).

**Figure 7:**
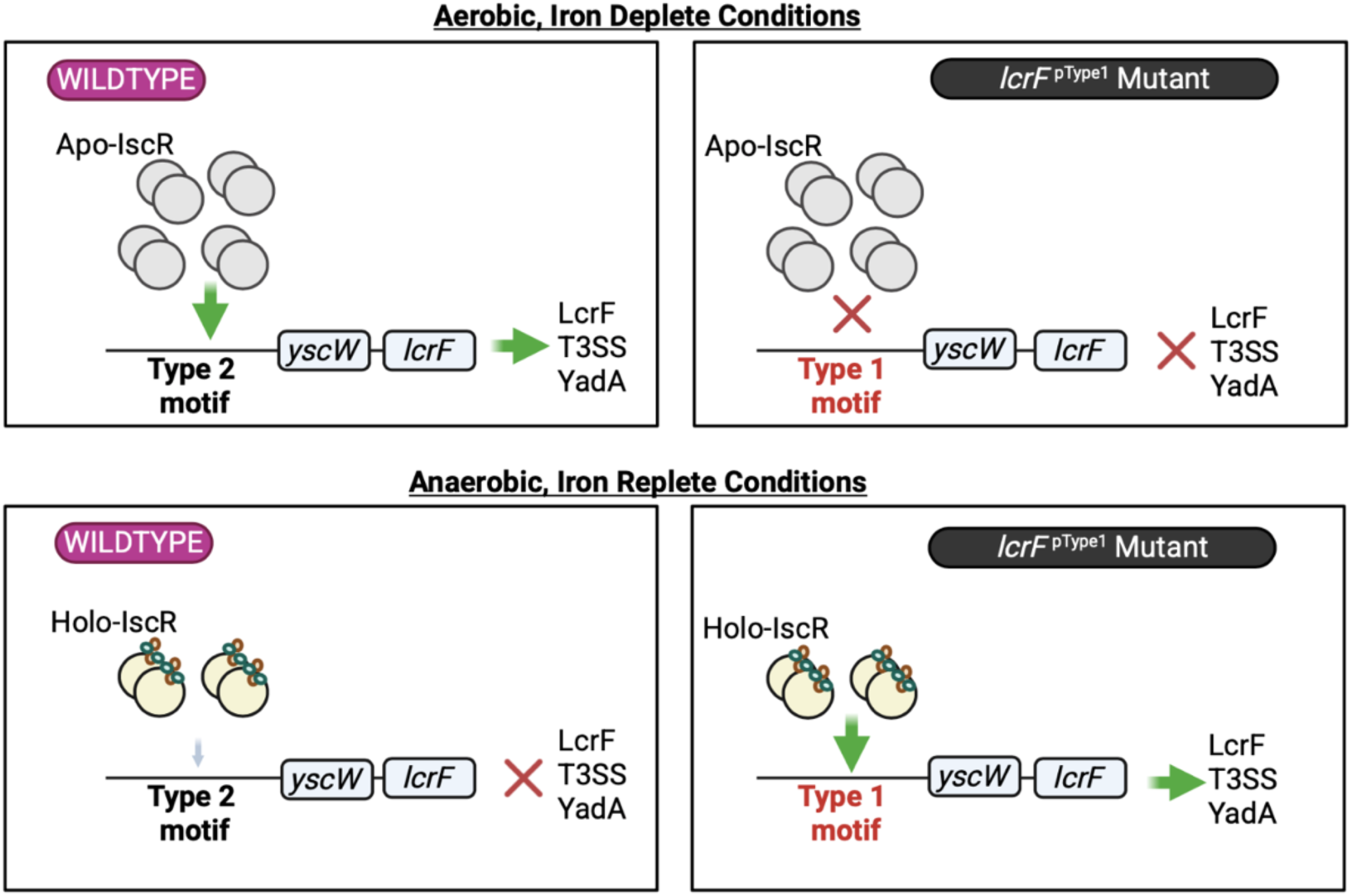
Schematic Representation of *lcrF* regulation by IscR in response to iron and oxygen. Wildtype *Y. pseudotuberculosis* encodes a *yscW-lcrF* promoter that contains a type 2 IscR binding motif. Apo-IscR levels are elevated under aerobic iron depleted conditions, where they are able to bind this type 2 IscR binding motif and promote *yscW-lcrF* transcription. The T3SS master regulator LcrF then drives transcription of the T3SS and the YadA adhesin. In contrast, holo-IscR is the predominant form of IscR under anaerobic, iron replete conditions and influences promoter activity through interaction with type 1, but not type 2, IscR binding motifs. Therefore, in wildtype *Yersinia*, LcrF, T3SS and YadA expression is repressed under anaerobic, iron replete conditions. In the pType1 *Y. pseudotuberculosis* mutant, the *yscW-lcrF* type 2 IscR binding motif has been swapped with a type 1 motif that only binds holo-IscR. In the pType1 mutant, LcrF, the T3SS, and YadA are ectopically expressed under anaerobic, iron replete conditions. Interestingly, the *Y. pseudotuberculosis* pType1 mutant experiences a tighter bottleneck to cecal colonization compared to the wildtype strain, suggesting that T3SS and/or YadA ectopic expression may be detrimental to *Yersinia* fitness in this anaerobic, iron replete environment.

Remarkably, we find that this simple switch yields a *Y. pseudotuberculosis* mutant with a T3SS expression profile opposite that of WT bacteria in terms of iron and oxygen responsiveness. We demonstrate that in anaerobic iron replete conditions, the *lcrF*^pType1^ mutant hyperexpresses LcrF, resulting in significantly greater T3SS activity and YadA adhesin expression relative to WT. Conversely, the *lcrF*^pType1^ mutant exhibits lower T3SS and YadA expression under aerobic iron limiting conditions, which normally induce robust T3SS and YadA expression in WT *Yersinia*.

Importantly, the *lcrF*^pType1^ mutant strain experiences a tighter bottleneck to colonization of the cecum, where *Yersinia* is predicted to experience an anaerobic iron replete environment. These data suggest that spatiotemporal sensing of iron availability and oxygen tension in host tissues by IscR allows *Yersinia* to deploy virulence factors in a strategic manner that optimizes bacterial survival.

Holo- and apo-IscR are both capable of binding to IscR type 2 binding motifs; however; holo-IscR may not influence transcription when recruited to a type 2 motif ^33^. In contrast, type 1 motifs are bound only by holo-IscR, since Fe-S cluster ligation by IscR is predicted to reorient a carboxyl group in the Glu43 residue that sterically prevents apo-IscR from binding type 1 motifs^31^. A recent study further showed that holo-IscR oxidized by O_2_ or redox cycling agents cannot bind to type 1 motifs or influence transcription from a type 2 motif^37^. *Y. pseudotuberculosis* and *Y. pestis* natively contain a type 2 IscR binding motif in the *yscW-lcrF* promoter, and our results show that changing this site to a type 1 motif prevents apo-IscR from binding, in agreement with previous biochemical studies. Because Fe-S cluster demand controls the levels of holo- and apo-IscR, switching the IscR binding motif in the *yscW-lcrF* promoter swapped the way in which iron availability and oxygen tension regulated expression of LcrF and its targets, YadA and the T3SS.

The <u>h</u>istone-like <u>n</u>ucleoid <u>s</u>tructuring protein (H-NS) and YmoA are known negative regulators of the *yscW-lcrF* promoter^24,48^. Previous work demonstrated that IscR acts as a roadblock to the repressive activity of H-NS/YmoA at the *yscW-lcrF* promoter^24^. It is possible that WT *Y. pseudotuberculosis* is unable to mount robust T3SS expression under anaerobic environments because holo-IscR cannot counter-silence H-NS/YmoA repression of the native *yscW-lcrF* promoter. In contrast, holo-IscR binding to the type 1 motif in the *lcrF*^pType1^ mutant *yscW-lcrF* promoter may allow counter-silencing of H-NS/YmoA under anaerobic conditions. Future work will explore the exact interactions of these proteins and their impact on T3SS activity and virulence in the host.

Switching the IscR binding motif in the *yscW-lcrF* promoter to reverse how iron and oxygen control *Yersinia* virulence factor expression led to altered infection dynamics. Previous data suggest that *Yersinia* encounters changes in iron availability and oxygen tension during the course of mammalian infection. *Y. enterocolitica* experiences iron replete conditions in the intestinal lumen as demonstrated by downregulation of iron and heme uptake pathways in a mouse model of infection^49^. Additionally, studies have shown that the healthy distal small intestinal lumen, cecum, and colon are predominantly low oxygen environments, sometimes described to be in a state of physiological hypoxia^40^. Building off these findings, we propose a model in which the *Y. pseudotuberculosis lcrF*^pType1^ mutant experiences a tighter colonization bottleneck in the anaerobic and iron replete conditions of the cecum. Based on our *in vitro* studies, an anaerobic iron replete cecal environment likely induces strong LcrF expression in the *lcrF*^pType1^ mutant but very little in the WT strain. These findings suggest that the T3SS and/or YadA hyper-expression in the cecum may be unfavorable for *Yersinia* cecal colonization. T3SS expression is metabolically burdensome and can lead to growth arrest in *Y. pseudotuberculosis*; perhaps this provides a disadvantage in the *lcrF*^pType1^ mutant if it is unable to grow to sufficient levels and fails to gain a foothold in the competitive niche of the cecal lumen.

Additionally, YadA hyperexpression in the intestinal lumen may lead to unfavorable adhesion to intestinal epithelial cells or autoaggregation^50, 46^. Lastly, it is possible that *lcrF*^pType1^ hyperexpression of the T3SS and/or YadA in the cecum may trigger an immune response that is detrimental to *Yersinia* fitness.

While *Y. pestis* has undergone substantial genome decay since its evolution from *Y. pseudotuberculosis*, *Y. pestis* has retained the IscR type 2 binding motif in the *yscW- lcrF* promoter^51^. While *Y. pestis* and *Y. pseudotuberculosis* have very different routes of transmission and infection cycles, IscR is required for T3SS expression in both species^38^. This suggests that the ability to promote LcrF expression specifically under conditions that promote high Fe-S cluster demand is beneficial for both *Y. pestis* and *Y. pseudotuberculosis*. Interestingly, IscR also regulates the *Salmonella enterica* SPI-1 T3SS but in response to conditions that promote low Fe-S cluster demand^52^. Whereas the *Yersinia* Ysc T3SS is important for colonization of extraintestinal tissues and may be detrimental to express in the cecum, the SPI-1 T3SS is important for *Salmonella* invasion of intestinal epithelial cells ^26, 53^. HilD is the master regulator of SPI-1 T3SS, and is autoregulated by binding to its own promoter ^54^ . In the *hilD* promoter, two IscR type 2 IscR binding motifs flank the HilD binding site, and a working model suggests IscR binding may outcompete HilD binding, although this has not been tested ^52^. Consistent with this notion, iron limitation leads to decreased SPI-1 T3SS expression, suggesting that IscR may allow *Salmonella* to sense Fe-S cluster demand as a spatiotemporal cue for crossing the intestinal barrier and downregulating the SPI- 1 T3SS when it is no longer needed. It is noteworthy that IscR was coopted in both *Yersinia* and *Salmonella* to couple sensing of Fe-S cluster demand to T3SS expression.

## Materials and Methods

### Bacterial culture conditions

*Y. pseudotuberculosis* strains used in this study are listed in Table 1. For anaerobic iron replete culturing conditions (Figures 1 and 3B), single colonies were inoculated into M9 media supplemented with casamino acids (referred to as M9 media hereafter) containing 0.2% glucose, and incubated aerobically at 26°C with shaking at 250 rpm. The following morning, the overnight cultures were subcultured to an OD_600_ of 0.1 into M9 with 0.9% glucose, introduced into the anaerobic chamber, and incubated at room temperature for 1 hour. Cultures were then transferred to the 37° C incubator in the anaerobic chamber and grown for an additional 5 hours.

**Table 1:** Strains.

| *Y. pseudotuberculosis* IP2666 Strain | Description | Source |
| --- | --- | --- |
| WT | Naturally lacks full-length YopT | <sup>55</sup> |
| *lcrF*<sup>pType1</sup> | IscR binding site in *yscW*-*lcrF* promoter mutated to a type1 binding motif | This work |
| $\Delta lcrF$ | In frame deletion of *lcrF* gene | <sup>56</sup> |
| *lcrF*<sup>pNull</sup> | IscR binding site in *yscW*-*lcrF* promoter mutated to abrogate any IscR binding | <sup>26</sup> |
| $\Delta iscR$ | In frame deletion of *iscR* gene | <sup>25</sup> |

For aerobic, iron starved culturing conditions (Figures 2 and 3A), cultures were grown as described ^26, 58^. Briefly, overnight cultures were grown at 26°C in M9+0.2% glucose, subcultured to an OD_600_ of 0.1 into chelated M9+0.9% glucose for 8 hours at 26°C, then subcultured again to an OD_600_ of 0.1 into chelated M9+0.9%glucose and incubated at 26°C for 12 hours. Finally, samples were subcultured an OD_600_ of 0.2 into chelated M9+0.9% glucose supplemented with 0.01 mg/L of FeSO_4_*7H_2_O, cultured for 2 hours at 26°C, and then for 4 hours at 37°C. All incubation steps were carried out aerobically with shaking at 250 rpm.

For culturing conditions depicted in Figure 4, overnight cultures grown in M9+0.2%glucose were subcultured into 3mL of M9+0.9%glucose into an OD_600_ of 0.2. The aerobic cultures were grown at 26°C for 1 hr, then switched to 37°C for 1 hr, both at 250RPM. The anaerobic cultures were placed in an anaerobic chamber at room temperature for 1 hr, then switched to 37 °C to continue anaerobic culturing for 2 hrs.

### Electrophoretic mobility shift assay (EMSA)

Untagged holo-IscR and tagged apo- IscR (IscR-C92A-His_6_) were purified as previously described^29,32^. PCR amplification of the wild-type *yscW*-*lcrF* promoter region (-206 to +12 bp relative to the +1 transcription start site) and its derivatives were carried out using primers XhoI-*lcrF* and BamHI-*lcrF* (Table 2) and plasmid templates in which the promoter of interest was cloned into pPK7179^59^ at the XhoI and BamHI restriction sites. The amplified DNA fragments were purified using the QIAquick PCR purification kit (Qiagen). DNA fragments (∼4 to 8 nM) were incubated with or without purified IscR-C92A- His_6_ (apo-IscR) as previously described^38^, except that after incubation at 37°C for 30 min, heparin was added to 50 µg/ml, and the mixture was further incubated at room temperature for 2 min before being loaded onto a nondenaturing 6% polyacrylamide gel. Gel electrophoresis was performed in 0.5X Tris-borate-EDTA (TBE) buffer at 200 V for 3.5 h at room temperature. Assays with holo-IscR were carried out in the same manner except the binding reactions and gel electrophoresis were carried out in a Coy anaerobic chamber, and gel electrophoresis was performed in 0.5X Tris-borate (TB) buffer with no EDTA. Gels were stained with SYBR green EMSA nucleic acid gel stain (Molecular Probes) and visualized using a Typhoon FLA 900 imager (GE).

**Table 2:**
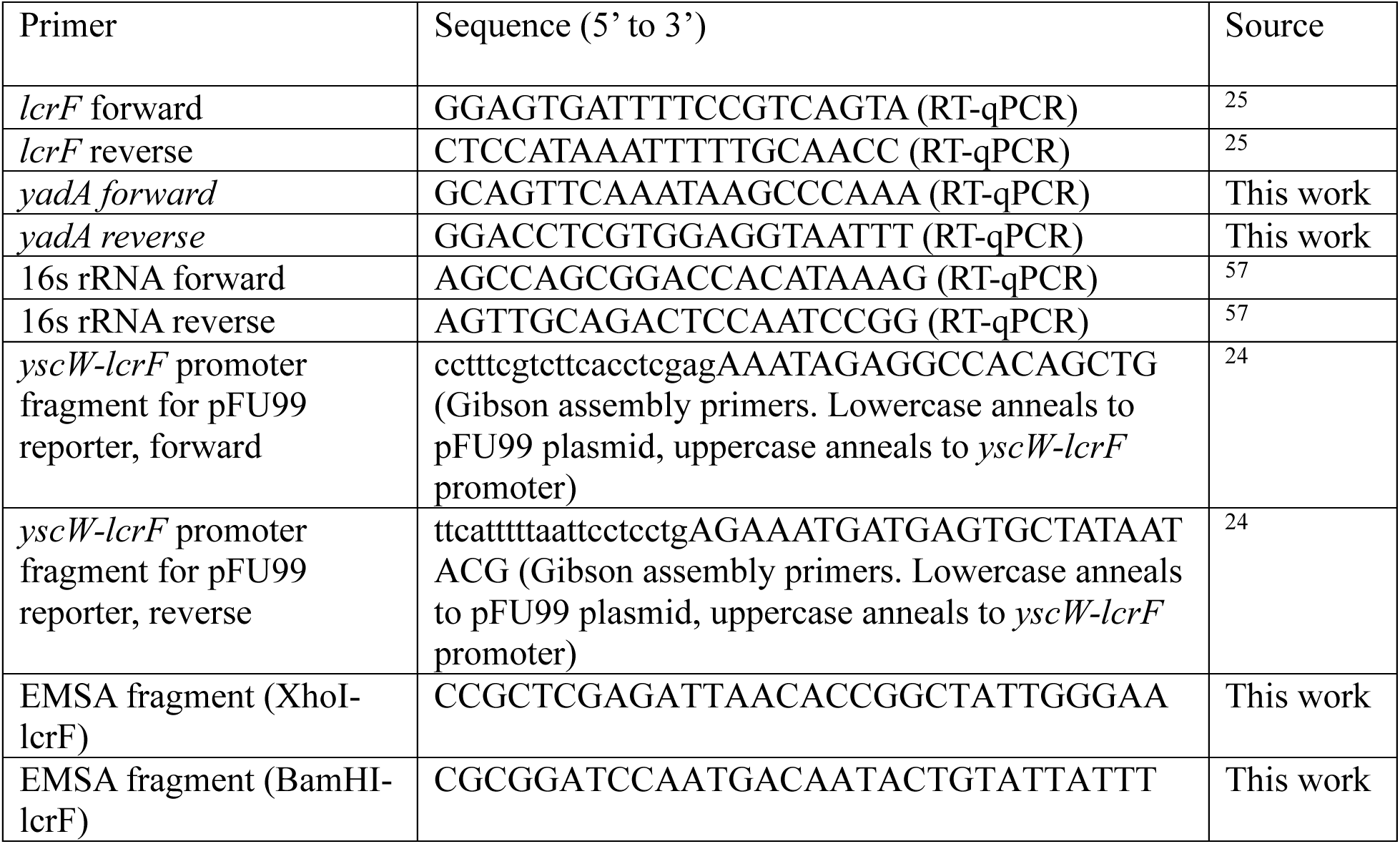
Primers.

### β-galactosidase Assay

pFU99 plasmids containing a chloramphenicol resistance cassette harboring either the wildtype or pType1 mutant *yscW-lcrF* promoter upstream of the lacZ gene were electroporated into both WT and Δ*iscR Y. pseudotuberculosis* strains. Samples were cultured either in anaerobic/iron replete or aerobic/iron starved conditions as described above, with the addition of 20 µg/mL chloramphenicol. Once the final incubation was complete, 5 µg/mL tetracycline was added, and samples were placed on ice for 10 min to halt further protein synthesis. A volume of 1 mL of each sample was centrifuged for 10 min at 3,381xg at 4°C and the supernatant was aspirated. Pellets were resuspended in 1 mL of ice-cold Z buffer, and the OD_600_ was recorded. One drop of 0.1% SDS and two drops of chloroform were added to each sample with a Pasteur pipette, and vortexed 1 minute to lyse the cells. 200 µl of 4 mg/mL ONPG was added to each sample, the time was recorded, and the sample was incubated in a 28°C water bath. Once a light-yellow color was observed, 500 µL of 1 M sodium bicarbonate was added to halt the reaction and the time was recorded. Samples were centrifuged for 1 minute at room temperature at 21,130xg, and 1 mL of each supernatant was transferred to cuvette. The OD_420_ and OD_500_ were measured, and Miller Units were calculated according to the following equation: MU=[1000*(OD_420_-1.75*OD_500_)]/(T*V*OD_600_), where T is the duration of the reaction in min, V is the volume of the final Z-buffer resuspended pellet that was added to each reaction.

### Western Blotting

Cultures were normalized to 1.8 mL of the lowest cell density culture and centrifuged for 15 minute at 21,130xg. Pellets were resuspended in 50 µl of final solution buffer (FSB) with DTT, heated at 95° C for 10 min, and stored at -70° C until use. Immediately prior to running on a gel, samples were heated again, centrifuged for 1 min at 21,130xg on a benchtop centrifuged, vortexed briefly, and 15 µl of each sample was run on a 12.5% SDS-PAGE gel, and wet-transferred onto a PVDF 0.45 µm transfer membrane. Blocking buffer contained 5% nonfat dry milk resuspended in TBST (Tris-buffered saline containing 0.1% Triton-X 100). All antibodies were diluted in blocking buffer. Briefly, membranes were blocked for 1 hour on a rocker at room temperature, then incubated with primary antibody (rabbit anti-RpoA was a gift from Melanie Marketon, rabbit anti-LcrF was a gift from Gregory Plano, rabbit anti-IscR^29^, or goat anti-YopE from Santa Cruz Biotechnology) overnight on a rocker at 4° C. Membranes were then washed in 10 mL TBST on a rocker for 5 min, for a total of three washes at room temperature. A 1:3000 titration of secondary mouse anti-rabbit or anti-goat IgG antibody conjugated to horseradish peroxidase (Santa Cruz Biotechnology) suspended in blocking buffer was then added, and membranes incubated for 1 hour on a rocker at 4° C. Finally, membranes were washed in TBST on a rocker for 5 min, for a total of three washes at room temperature. For imaging, 2.5 mL of Solution A and 2.5 mL of Solution B from the luminol reagent kit (Santa Cruz Biotechnology) were added to the membrane, incubated for 1 minute with manual vigorous shaking, and the BioRad ChemiDoc imaging system was used to visualize the membranes. ImageLab software was used to calculate relative protein quantification. RpoA, LcrF, IscR, and YopE were also normalized for each sample relative to WT levels. Relative expression levels were calculated by dividing the normalized values of the protein of interest by the normalized values of their respective RpoA loading control value.

### T3SS activity assay

Cultures were normalized to 1.8 mL of the lowest cell density culture and centrifuged for 15 min at 21,130xg . Culture supernatants were filtered using a Millex-GV PVDF 0.22um filter attached to a syringe. 10% of the supernatant volume of 6.1 N trichloroacetic acid (TCA) was added to each sample, in addition to 4 µl of 0.5 mg/mL BSA as a protein precipitation control. Samples were briefly vortexed and placed in ice in a 4° C room overnight. To collected precipitated protein, samples were centrifuged at 21,130xg at 4° C for 15 min and supernatants were aspirated. To remove residual TCA, 1 mL of ice-cold acetone was added to each tube (without resuspending), centrifuged at 21,130xg for 15 min at 4°C, and the acetone was carefully aspirated. The acetone washing step was repeated once more and pellets were left to dry at room temperature for approximately 30 min. Pellets were resuspended in 50 µl FSB with DTT, heated for 10 min at 95°C and stored in -70°C until use. Immediately prior to running on a gel, samples were heated again, centrifuged for 1 min at max speed on a benchtop centrifuged, vortexed briefly and 5 µl was loaded into a 12.5% SDS-PAGE gel. Gels were then incubated in Coomassie blue stain for 2 hours at room temperature on a rocker, then destained overnight at room temperature until all background color was removed (approximately 2-3 changes of destain were necessary to achieve proper contrast between gel and protein bands). Gels were imaged using the BioRad ChemiDoc and ImageLab software.

### RNA isolation and quantitative PCR (qPCR)

A total volume of 5 mL of culture for each sample was pelleted for 5 min at 3,968xg and pellets resuspended in 500 µl of the same media used for culturing the cells. A total volume of 1mL RNA Protect reagent (QIANGEN) was added and RNA was isolated according to the manufacturers protocol using the RNeasy Mini Kit (QIAGEN). A Turbo DNA-free Kit treatment (ThermoFisher) was used to remove DNA contamination according to the manufacturer’s protocol. cDNA was generated using the M-MLV (ThermoFisher) Reverse Transcriptase according to the manufacture’s protocol. Each 15 µL PCR reaction contained PowerUp SYBR Green Master Mix (ThermoFisher), primers (Table 1), and either 1:10 dilution of cDNA for the gene of interest or 1:1000 dilution of cDNA for 16s rRNA. The 16s rRNA expression levels of each sample were used to normalize gene of interest expression levels. Results for each strain are represented as fold change relative to WT.

### Generating STAMPR libraries

Briefly, the wildtype and *lcrF*^pType1^*Y. pseudotuberculosis* IP2666 strains were used to create the barcoded libraries. The diaminopimelic acid (DAP) auxotroph MFDλpir pSM1 *E.coli* library containing the Tn7 transposon harboring a unique ∼25bp barcode with a kanamycin resistance cassette was used as the donor library which is known to integrate in a neutral site downstream of the *glmS* gene in the recipient chromosome^60^. A diaminopimelic acid auxotroph MFDλpir pJMP1039 *E. coli* strain was used as a helper strain to carry out the triparental mating. 30µl straight from the glycerol stock of the donor pSM1library was used to start a 3mL LB+DAP(300µl) +Kan (50µg/mL)+Carb(50μg/mL) overnight culture at 37°C at 250 RPM; a single colony of the helper pJMP1039 strain was used to start a 2mL LB+ DAP(300µl)+Carb (50 μg/mL) overnight at 37°C at 250RPM; and a single colony of the appropriate recipient *Y. pseudotuberculosis* strain was used to start a 2mL LB overnight culture in 26°C at 250RPM. 100μl of overnight cultures of the donor library, the helper strain, and the *Yersinia pseudotuberculosis* recipient strains were mixed in equal parts, centrifuged at 3,381xg for 5 min, then resuspended in 100µl of PBS and the mating was carried out for 24 hr. Mating was done by placing a 0.45um MCE membrane (25mm) at room temperature on a 10cm LB+DAP(300µM) plate, and pipetting 100µl of the mixed strains on top; a total of 6 filter papers were used on a single 10cm plate, each containing 100µl of mating mixture. After the mating was complete, cells from all 6 filter papers were pooled into 1mL of PBS and 900µl of this mixture was plated on a 15cm LB+Kan (50µg/mL) plate and incubated for 48 hr at room temperature, the rest of the 100µl was used to plate dilutions to ensure proper transconjugant efficiency. Transconjugants were scraped, resuspended in LB+25% glycerol and aliquots were stored in -80° C.

### Mouse experiments

All experiments were approved by the University of California Santa Cruz Institutional Animal Care and Use Committee and were performed in strict accordance with National Institutes of Health guidelines. Nine-week old female C57BL/6J mice were obtained from Jackson Laboratory (Strain #000664). Food was withheld from mice for approximately 16 hours prior to infection. Briefly, 30 µL of the glycerol stock for each *Y. pseudotuberculosis* STAMPR library was cultured overnight at 26° C in 3 mL of LB supplemented with 50 µg/mL of kanamycin.

Appropriate volumes of the cultures were pelleted at 3,381x g for 5 min at room temperature. The pellets were resuspended in PBS to obtain 1x10^9^ CFU/5 µL inoculum. A total of 5 µl of the inoculum was placed on a small piece of mouse chow and introduced to mice for consumption, then food and water were provided *at libitum.* Four days post-inoculation, organs of interest were harvested and homogenized. A portion of the homogenate was used to plate serial dilutions on 10 cm LB agar plates containing 2 µg/mL irgasan and 50 µg/mL kanamycin to determine CFU/g tissue. The entirety of the remaining homogenate was plated on a 15 cm LB agar plate containing 2 µg/mL irgasan and 50 µg/mL kanamycin for STAMPR analysis. Plates were left at room temperature to grow for 48 hours. A total of 1mL of PBS was used to scrape each STAMPR plate and 10 µl of sample was added to 90 µl of water, boiled for 15 min at 95° C in a thermal cycler, and processed for sequencing and previously described ^61,62,63^.

### Myeloperoxidase (MPO) ELISA

Mouse fecal pellets were collected and weighed and resuspended in 1mL of PBS containing protease inhibitors (Roche cOmplete ULTRA Tablets). Samples were homogenized using glass beads prior to storage in - 70 °C until use. Thawed samples were centrifuged at 400xg and 100ul of each sample was used to determine abudnace of MPO using the mouse myeloperoxidase DuoSet ELISA kit (R&D Systems).

### Read Processing and Calculating Ns

Briefly, the raw output of reads were obtained from an Illumina NextSeq 1000 instrument. Samples are multiplexed in a 8x8 grid where each row contains a distinct forward primer and each column contains a distinct reverse primer. Each sample thus contains a distinct pair of primers (combinatorial indexing). Demultiplexing via the column is performed using i7 while demultiplexing of the row is performed with the first 12 nucleotides of the read. Demultiplexed reads were trimmed and mapped to the corresponding pSM1 reference^64^. The “preprocessReads” function from the QuasR R package was used to trim reads, and the “qAlign” function from the QuasR R package was used to map the trimmed reads. The mapped reads were then corrected for index hopping as described previously by Holmes et al.^65^, and Ns was calculated as described by Hullahalli et al.^47^. Briefly, the reference (inoculum) vector was iteratively resampled through simulated bottlenecks across a range of sampling depths (founding population sizes), starting from 1 read to 20 times the total number of barcodes in the library. A plot was generated that mapped the simulated sampling depth (x-axis) to the number of unique non-zero barcodes (y-axis) detected in each simulation. Ns represents the sampling depth that yields the observed number of unique barcodes in a sample, determined by inverse interpolation from this graph.

## Acknowledgements

This study was supported by National Institutes of Health (www.NIH.gov) grants R01AI119082 (to V.A. and P.J.K.) and RO1 AI-042347 (to M.K.W.) as well as the Howard Hughes Medical Institute (M.K.W.). M.O. received support from the UC Santa Cruz Chancellor’s Fellowship. The funders had no role in study design, data collection and analysis, decision to publish, or preparation of the manuscript. Figure 7 graphical image created using BioRender.com.

V.A. and M.O. conceived of the study and devised the experiments. M.O., G.I.B., D.F.P., C.L., K.G.D, E.M. D.H.R., and L.S. performed the experiments. M.O., G.I.B., K.H., E.M., and V.A. analyzed the data. P.J.K., M.K.W., and V.A. acquired funding and provided supervision. M.O. and V.A. wrote the original draft and M.O., G.I.B., K.H., E.M., P.J.K., M.K.W., and V.A. reviewed and edited the manuscript.

## Supplemental Figures

**Supplemental Figure 1:**
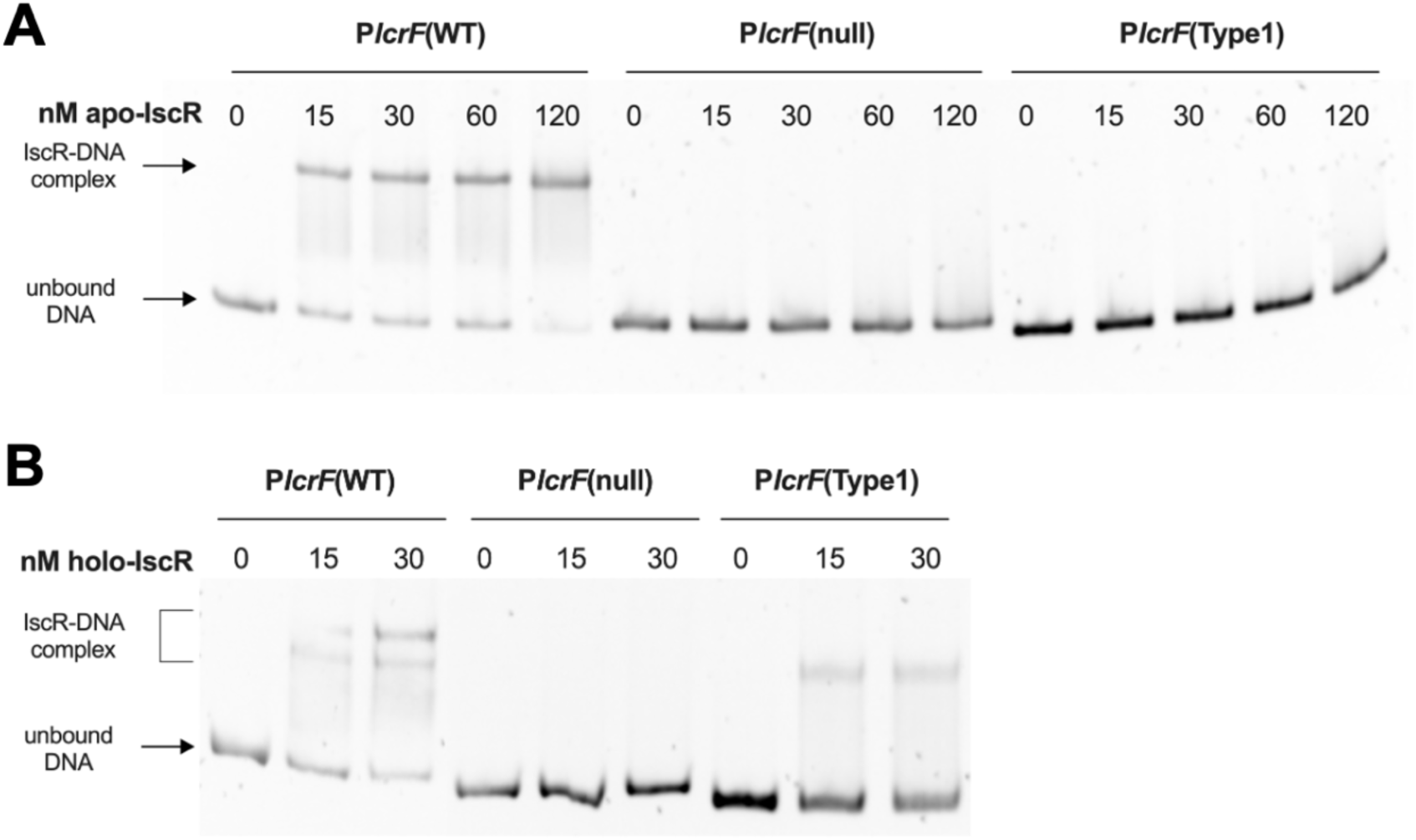
Holo-IscR, but not apo-IscR, binds to the *lcrF*^pType1^ mutant promoter. **A)** Electrophoretic mobility shift assay (EMSA) with increasing concentrations of IscR-C92A (apo-IscR) and either the WT *yscW-lcrF* promoter or the *lcrF*^pType1^ mutant promoter. **B)** EMSA carried out under anaerobic conditions with increasing concentrations of IscR prepared under anaerobic conditions (holo-IscR) and either the WT *yscW-lcrF* promoter or the *lcrF*^pType1^ mutant promoter.

**Supplemental Figure 2.**
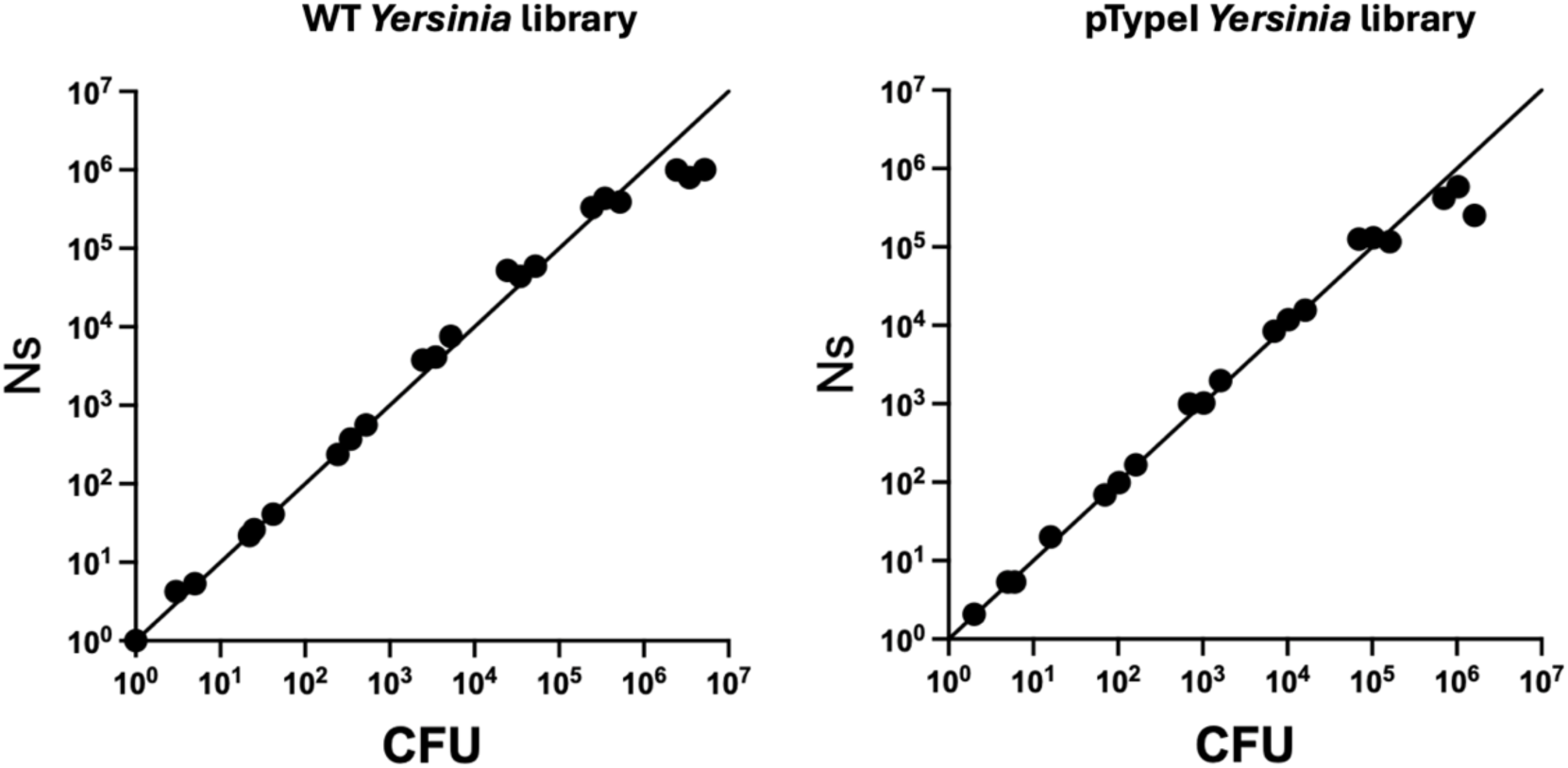
Y*e*rsinia STAMPR *in vitro* bottleneck. STAMPR libraries were made in the WT and *lcrF*^pType1^ *Y. pseudotuberculosis* backgrounds. Bacteria were grown in LB at 26°C and serial dilutions plated onto LB agar. Bacterial mass was collected from each plate and barcodes sequenced. The x-axis represents the known founding population (FP) size following culture dilution (*in vitro* bottlenecks) and the y-axis represents the calculated founding population from barcode sequencing (Ns). The high correlation between Ns and CFU demonstrates that both libraries accurately quantify bottlenecks up to FP = 5 x 10^5^, consistent with previous STAMPR libraries.

